# Megacolonies: an alternative social organization in anemonefishes?

**DOI:** 10.1101/2022.11.06.515354

**Authors:** Manon Mercader, Jann Zwahlen, Kina Hayashi, Hiroki Takamiyagi, Yung-Che Tseng, Hai-Thanh T. Nguyen, Keishu Asada, Jérome Sowinski, James Reimer, David Lecchini, Vincent Laudet

## Abstract

Anemonefish are iconic examples of marine fishes living in mutualistic symbiosis with sea anemones. In a given sea anemone, the anemonefishes have a stereotyped social organization with a dominant female, a semi-dominant male, and several juveniles. A strict size-based hierarchy governs the social interactions within these colonies, with each individual differing from the previous or next fish in the order by +/- 20% size. This social organization is conserved across the Indo-Pacific in all 28 species of anemonefish found on any of ten giant sea anemone species. We report the existence of huge “megacolonies” of up to 100 fish living in large carpets of sea anemones. This alternative organization was observed for different fish and anemone species in different coral reef locations (French Polynesia, Japan, Taiwan, and Vietnam). In these colonies, the strict size-based hierarchy is no longer recognizable, and the level of aggressivity of the different members appears lower than in “normal” colonies. These megacolonies may correspond to a previously overlooked type of social organization that could be linked to host availability and offer a unique opportunity to understand anemonefish’s behavioral, social, and hormonal plasticity.

## Introduction

Many animal species live in societies displaying a wide range of possible organizations, from stable pairs, shoals of fish, flocks of birds, or swarms of insects to the eusocial organizations that exist in many insects and mammals [1–3]. Social groups organize themselves in myriad ways, and these organizations impact the whole life of animals, whether at the level of reproduction, food intake, or defense against predators. Studying these modes of organization, their functioning, their robustness, and also their flexibility and plasticity is essential to understand the organization of ecological systems and to realize how they can adapt to environmental changes and, in particular, to anthropogenic stresses [4,5]. Therefore, analyzing social organizations has allowed biologists to understand multiple facets of social organizations, from the mechanistic processes involved to the study of ecological and evolutionary functions [6].

The study of patterns of social organization in marine animals is particularly demanding due to both the difficulty of conducting long-term observations and the marine environment’s temporal and spatial dynamism [7]. Thanks to biologging approaches combined with intensive observation programs, we now have a much better idea of the social interactions between individuals in many groups of cetaceans and fish, but work until now has generally been limited to comparatively large animals [8,9]. Although there are many observations suggesting social organizations and elaborate behaviors for other species of marine fish, we are still far from the same understanding of social organizations as has been achieved for many terrestrial animals [7]. In coral reefs, fish are widely studied regarding their ecology, behavior, and social organization [10]. Within this ecosystem, anemonefish form a group that has been particularly studied from this point of view.

In this clade of 28 species of Pomacentridae, scientists have developed a preliminary integrated understanding of social organization from ecological to molecular levels (reviewed in Laudet and Ravasi, 2022 [11]). Indeed, these fish, which live in mutualistic symbiosis with giant sea anemones, never abandon their host sea anemone and therefore form elaborate micro-societies that can relatively easy be studied at sea [10–12].

The social structure of anemonefish within their host anemone is highly organized. This organization consists of a hierarchy based on size: within a colony, no individual of the same size exists, and the different fish are classified in descending order of size, with an average difference of 20% between each rank [13,14]. At the top of the hierarchy, a dominant female will aggressively defend the colony and maintain her ascendancy over the smaller members. The second individual in size is the male who will reproduce with the female and take care of the eggs, aerating them, removing dead eggs, and of course, also defending them against possible predators. Thus, both parents exercise parental care to allow the proper development of the eggs [15]. Finally, the colony contains a variable number of sexually immature juveniles, again ranked by size, forming a queue waiting for access to reproduction. If the female dies, the male transforms into a female, the largest of the juveniles into a male, and each subsequent juvenile gains a place in the line [14,16]. Because of this social organization, anemonefish present exciting opportunities to generate new concepts and test the robustness of current theories of social evolution. Organized colonies of anemonefish thus raise many questions: Why juveniles give up their own reproduction for a very long time? Why do breeding adults tolerate juveniles within colonies? How are conflicts between colony members resolved? (See review by Buston et al., 2022, which discusses these different questions in detail[14]). Anemonefishes allow addressing all these questions, which are among the major objectives of behavioral ecologists and evolutionary biologists.

Many indications suggest that this social organization is conserved within all 28 species of anemonefish, which associate, in a non-random way, with ten species of giant sea anemones [12,17]. It is, however, still unclear if this organization can be plastic and, in particular, how changes in ecological constraints could eventually lead to different organizations. By observing colonies of different anemonefish species at different coral reef locations and under different ecological contexts, we have observed colonies that do not obviously fit with the precise organization presented above. In this paper, we describe “megacolonies”, in which the strict social hierarchy based on size does not seem to operate as rigidly as in “normal” cases. Such observations could provide fascinating opportunities to study the plasticity of social organizations when ecological constraints vary.

## Material and Methods

### Field observations

Observations of alternative colony structures were done while scuba diving or snorkeling during various surveys and sampling activities. When possible, the noted colonies were revisited to determine the colony’s structure (i.e., number of individuals (fish and anemone), fish social status, and presence of other species) via Underwater Visual Census (UVC) methods [18–20].

### Study sites

Bora-Bora, French Polynesia. One megacolony was observed in the lagoon of Bora-Bora (16°29’S, 151°44’W), French Polynesia; a volcanic island formed 3.45 to 3.10 million years ago in the tropical South Pacific. The coral reefs surrounding Bora-Bora have an area of about 70 km² [21]. Although there are several classic *A. chrysopterus* (the only anemonefish species present in French Polynesia) colonies in the area, the megacolony was discovered on a turbid sandy area in the barrier reef (16°26’59.07”S; 151°44’44.46”W) in 2021 and has been monitored since.

Kagoshima, Japan. This location hosts high densities of *A. clarkii* (mainly living in *E. quadricolor*), the only anemonefish species present in mainland Japan. Most of them live in “normal” colonies, but several megacolonies were observed in July 2022 in Kagoshima Bay (31°22’N, 130°40’E) on the southern coast of Kyushu (East China Sea). This long (about 60 km from the end to the mouth of the bay) and enclosed bay is partially of volcanic origin, and two submarine calderas mainly shape its shoreline, Aira Caldera in the north and Ata Caldera at the southern mouth, and formed 22,000 and 150,000 years ago, respectively. The bay’s northern end hosts large yellowtail (*Seriola quinqueradiata*) and amberjack (*Seriola dumerili*) fish farming facilities, and the underwater substrate is mainly composed of rock and muddy bottoms, making the water often turbid. At the bay’s entrance, the bottom is mainly composed of rock and sand, and the water is clearer. Kagoshima Bay has a warm temperate climate (water temperature varies from an average of 16.5°C in winter to 28.5°C in summer).

Okinawa, Japan. Okinawa Island (26°28’N, 127°50’E) is part of the Ryukyu Archipelago in southern Japan. It has a humid subtropical climate. Despite its relatively high latitude, the water temperature varies from an average of 20°C in winter to 28°C in summer due to the northward flowing warm-water Kuroshio Current. The island is surrounded by highly diverse fringing and patch reefs, but the coast is also highly modified by land reclamation [22]. Okinawan waters are home to six species of anemonefish (*A. clarkii, A. frenatus, A. ocellaris, A. perideraion, A. polymnus*, and *A. sandaracinos*) living in association with seven anemone species [23]. Megacolonies were observed at several spots around the island; in Oura Bay on the east coast (26°33’5.48”N, 128°2’18.47”E), Atsuta Beach on the west coast (26°30’51.91”N, 127°53’45.00”E), and Chinen Peninsula in the south (26°10’14.17”N, 127°49’53.47”E), all between January and August 2022.

Nha Trang Bay and Van Phong Bay, Vietnam. While the coral reefs of Nha Trang Bay are well-known due to a long history of research (e.g., [24,25]). They face many anthropogenic pressures [26,27], as opposed to more pristine Van Phong Bay. Megacolonies were found on sand/rubble areas and were observed in both bays (Nha Trang Bay, 12°10’14.04”N, 109°18’43.20”E; Van Phong Bay, 12°34’15.24”N, 109°23’58.30”E) during surveys in July 2022, with field notes taken along with videos and images. Water temperatures during the surveys were 27°C to 30°C. Kueishan Island, Taiwan. Kueishan Island is located northeast of Taiwan (24°50’N, 121°57’E). The island is a geologically young and active volcanic island in Taiwan, and its hydrothermal vents create a unique ecosystem around the island [28–30]. Water temperature varies between 20°C and 28°C in the non-vent areas [31], and patchy coral communities surround the island. The megacolony was observed on a rocky bottom with a few corals at the island’s eastern tip (24°50’29.9”N, 121°56’17.1”E).

## Results

As described above, colonies with organizations deviating from the strict social structure of “normal” colonies were found in different geographical locations and environmental conditions. They involved various species of anemonefish and anemones (Fig 1 and Table 1). These different types of alternative organization were classified into two categories: intraspecific megacolonies (i.e., composed of a large number of anemonefishes of the same species living in a large number of host anemones of the same species) and interspecific megacolonies (i.e., composed of a large number of anemonefishes from several species living in various species of host anemones) (Fig 2). Detailed examples of each type of megacolony are given below.

**Figure 1:**
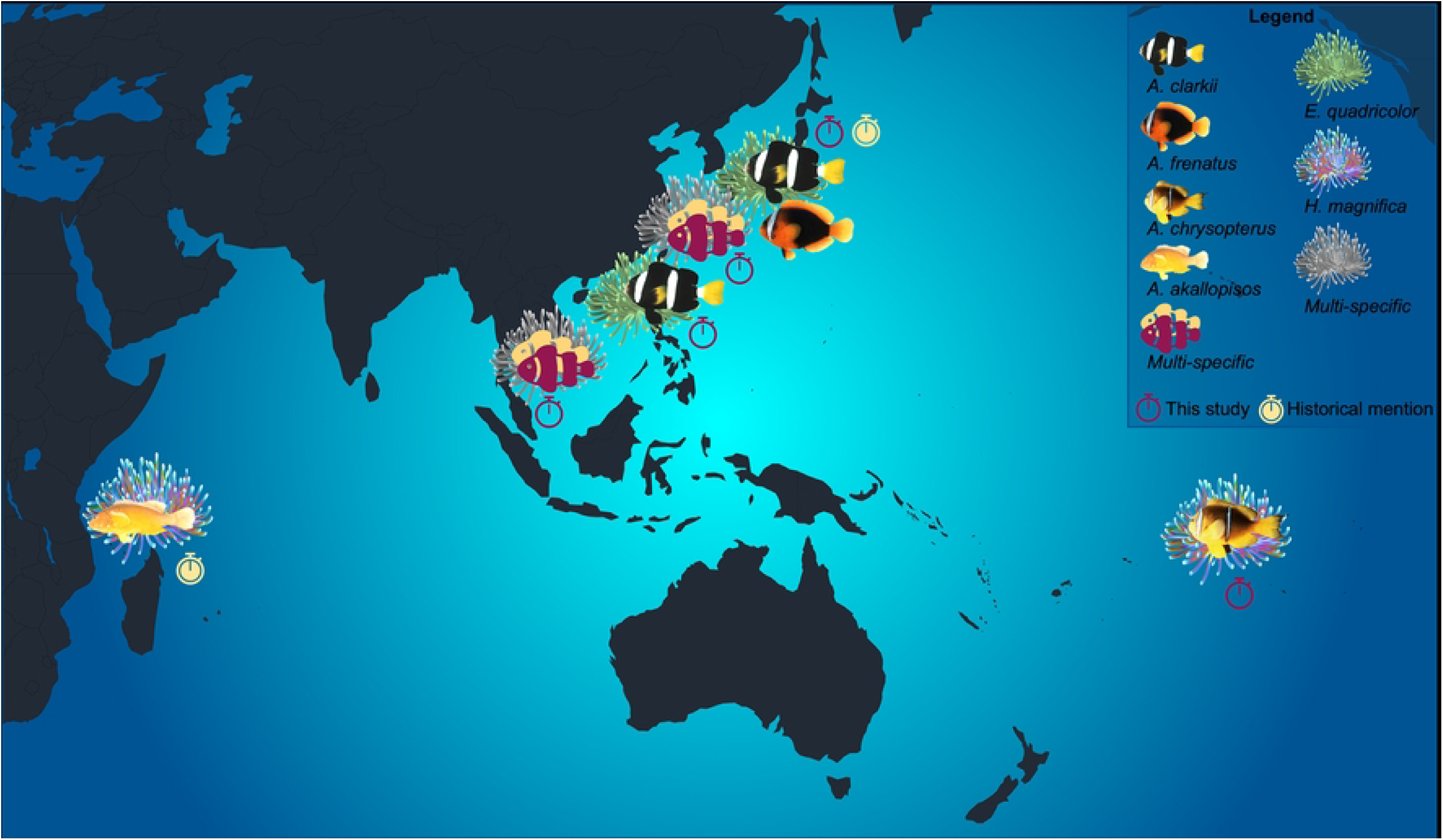
Geographical location of the two types of megacolonies described in this study and in previous studies with their fish and anemone species compositions.

**Table 1:**
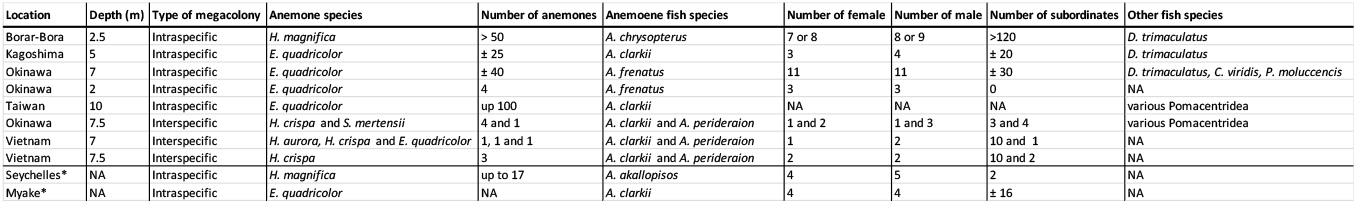
Summary of the different megacolony types. * indicates mentions in the scientific literature.

**Figure 2:**
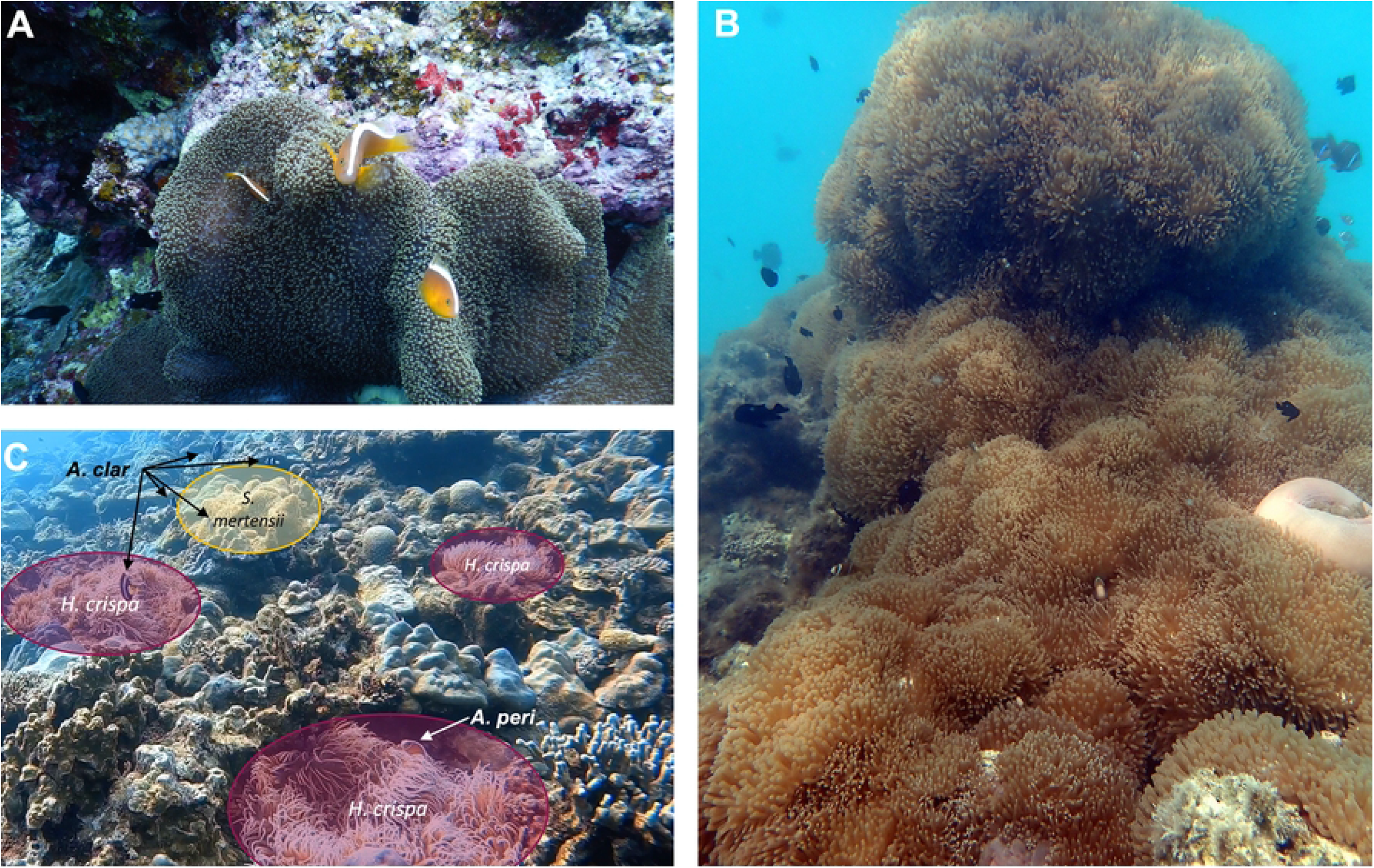
(A) A “normal” colony (female, male and one juvenile) of *A. sandaracinos* on *S. mertensii*, (B) intraspecific megacolony in Bora-Bora, French Polynesia, composed of an *H. magnifica* carpet with several *A. chrisopterus* and *D. trimaculatus* (more images in S1 Fig.) and, (C) interspecific colony in Atsuta beach, Okinawa, Japan, in the picture all *A. clarkii* and one *A. perideraion* are visible as well as all 5 anemones (left *H. crispa* is actually two individuals with overlapping tentacles) (additional images in S4 Fig. and S5 Fig).

### Intraspecific megacolonies

Bora-Bora, French Polynesia. This megacolony was composed of a carpet of the anemone *Heteractis magnifica* on which the anemonefish *A. chrysopterus* lived. More than 50 *H. magnifica* covered up to 95% of a dead coral patch of 3 m in length and 2 m wide (Fig 2-B and S1 Fig) at a depth of 2.5 m. The anemonefish population was estimated to be comprised of seven or eight females, eight or nine males, and more than 120 sub-adults and juveniles. When scared by a diver, adult fish swam around and hid in the anemones but always returned to a well-defined site within the megacolony. In this megacolony, the anemonefishes lived with a large three-spot *Dascyllus* population (more than 150 individuals of *Dascyllus trimaculatus*).

Sata, Kagoshima, Japan. This megacolony was composed of many *A. clarkii* living on a carpet of the anemone *Entacmea quadricolor*. The site was a rocky bottom, about 5m deep, with more than 25 anemones mainly in cracks covering about 10% of an approximately three 3 × 5 m area (S2 Fig). The colony was composed of three breeding pairs, one additional male, and approximately 20 sub-adults and juveniles. Young recruits stayed within the tentacles of a specific anemone, while bigger immature individuals were observed swimming from one anemone to the other. Mature fish would also enter different anemones but always seemed to return to the same spot. Few aggressive interactions were observed between *A. clarkii* individuals. A few *D. trimaculatus* individuals were also found around this sparse anemone carpet. Several megacolonies of this type were observed in this geographical area, but only one was described in detail.

Oura Bay, Okinawajima Island, Okinawa, Japan. This megacolony was composed of a carpet of the anemone *E. quadricolor* upon which live *A. clarkii* anemonefish. The site is a dead coral patch (approximately 2 m in diameter) on a muddy bottom. The base is 7 m deep, and more than 40 anemones cover approximately 80% of the top part of the patch (around 5 m deep) (S3 Fig). The anemonefish population was estimated to be 11 breeding pairs and approximately 30 subadults and juveniles. The coral patch also hosted many *D. trimaculatus, Chromis viridis*, and *Pomacentrus moluccensis*. No behavioral data were collected.

Chinen Peninsula, Okinawajima Island, Okinawa, Japan. A smaller megacolony of *A. frenatus* in *E. quadricolor* was observed in the southeast of Okinawajima Island. It comprised four anemones hosting three breeding pairs (S3 Fig). *A. clarkii* and *A. clarkii* fish seemed to be swimming freely from one anemone to the other but eventually returned to the same host individual.

Kueishan Island, Taiwan. This megacolony was composed of a carpet of the anemone *E. quadricolor* upon which live *A. clarkii* anemonefish. The site is a 10m deep rocky bottom with some corals. Over a hundred *E. quadricolor* individuals cover an approximately 50m^2^ area. A large number of adults and juveniles inhabit this megacolony. However, no detailed estimation of the colony’s structure was performed. Various species of Pomacentridae (e.g., D. *trimaculatus*) and Labridae also live in this anemone carpet (S4 Fig).

### Interspecific megacolonies

Atsuta Beach, Okinawajima Island, Okinawa, Japan. This megacolony comprised four *Heteractis crispa* and one *Stychodactyla mertensii* within about 2m^2^ of a mix of dead and live scleractinian corals, with an anemone coverage of approximately 30%. The site was 7.5 m deep. *A. perideraion* inhabited the four *H. crispa*. One anemone hosted only one individual, another a colony composed of a breeding pair, and one large subadult. The last two anemones were next to each other and together hosted a colony consisting of a breeding pair and three juveniles. The *S. mertensii* was inhabited by an *A. sandaracinos* colony (breeding pair and one juvenile) and an *A*.*clarkii* colony (breeding pair and three juveniles) (Fig 2-C and S5 Fig). Aggressive interactions between *A. clarkii* juvenile and *A. sandaracinos* individuals were observed but not between adults (S6 Fig). Adult *A. clarkii* also entered the neighboring *H. crispa* without aggressive interactions with resident *A. perideraion* individuals (S6 Fig). A high density and diversity of damselfish (e.g., *P. lepidogenys, P. alexanderae, Pomachromis richardsoni, Chromis chrysura, Amblyglyphidodon curacao*), as well as several *Labroides dimidiadus* individuals, were observed around this megacolony (S5 Fig).

Van Phong Bay, Vietnam. This megacolony was composed of three different species of host anemone; *H. aurora* (size 22 × 22 cm), *H. crispa* (35 × 45 cm), and *E. quadricolor* (30×30 cm), all within 5 m of each other, at depths of 6.5 to 7.6 m on the north coast of Hon Lon Island, on rubble/sand substrate. The three anemones were inhabited by a large number of *A. clarkii* (three adults and ten juveniles), which aggressively defended all three anemones. The mature fish constantly swam between the three anemones, while juveniles remained with a single anemone (7 on *H. aurora*, three on *H. crispa*). We did not observe any aggressive behavior between *A. clarkii* individuals. One of the anemones (*H. crispa*) also contained a single *A. perideraion*.

Nha Trang Bay, Vietnam. This megacolony consisted of three *H. crispa* anemones (diameters 25 × 30 cm, 30 × 30 cm, 35 × 35 cm) within 3 m of each other at 7.3 to 8.1 m depth, within the marine protected area at Hon Mun, Nha Trang Bay. The three anemones were inhabited by a large number of *A. clarkii* (n=14, at least four adults, remainder juveniles), which aggressively defended all three anemones. As in Van Phong Bay, mature fish constantly swam between anemones, while juveniles remained with a single anemone (n= 5, 4, and 1, respectively). We did not observe any aggressive behavior between *A. clarkii* individuals. Two of the anemones also contained a single *A. perideraion*.

## Discussion

These megacolonies of anemonefish living in different host anemone species and geographical locations might be more common than previously thought. This situation raises new scientific questions, notably in terms of socio-evolution, while providing a model to address them.

In the past literature, we did not find any mention of the term “megacolony.” However, several older studies have described alternative social organizations in anemonefish under different names, such as “super anemones” [32] or “multi-adult social groups” [33]. We found also work from the 1970s reporting the existence of such megacolonies in different geographical locations and for various species (Fig 1 and Table 1). In Aldabra, Seychelles, carpets of up to 198 *H. magnifica* individuals have been observed. They were divided into groups of up to 17 individual anemones hosting as many as nine adult and several juvenile *A. akallopisos* [32]. In Miyake-jima, Japan, *A. clarkii* was reported to form groups of 20 to 24 fish (four breeding pairs) on a 14m^2^ area partially covered by *E. quadricolor* [33–36]. Ten years later, *A. clarkii* megacolonies from Shikoku, Japan, were used to investigate reproductive behavior and territory acquisition [37–40]. Thus, there appears to be a wealth of information from the 1970s to early 1990s, but, to our knowledge, there have been no recent studies on these types of colonies, as well as no previous descriptions of any inter-specific megacolonies. Below, we highlight some research avenues we believe megacolonies could help address.

### Plasticity in social structure and mating system

Numerous studies have described the anemonefish’s social structure as very stable and conserved (reviewed in [11]). However, our observations combined with those from past studies mentioned above suggest plasticity in anemonefish behavior and social organization. For instance, Fricke (1979) described a typical colony social structure within the *A. akallopisos* megacolony. In their investigated site, each female defended a territory of a maximum of 0.88 ± 0.12 m^2^. Exceeding this surface area, a female cannot protect her territory against competitors, which determines the spacing between colonies (breeding pairs). However, juveniles were swimming freely from one territory to another [32]. Moyer (1980) observed competition between *A. clarkii* breeding adults after the breeding season, which could lead to the “displacement” of some individuals by more competitive ones. Displaced adults lived in a coral near anemone patches and eventually displaced other adults to conquer a new territory and breeding position. Moyer (1980) also reported long-distance travel (over 50 m away from an anemone) and clustering behavior (i.e., several adults coming together about 20 m away from their anemones)[33], which is more reminiscent of the damselfish *Dascyllus aruanus’* social organization [41]. In the megacolonies we observed, fish were more mobile and seemed less prone to aggressiveness than in “normal” colonies. Social interactions and behavior of bigger social groups should be investigated in more detail, for example, using Social Network Analysis (SNA) [6,42,43]. Megacolonies could represent useful models to assess how social systems vary when ecological constraints change (in this case, change in habitat availability) and test several theories in social evolution [10].

Buston (2022) and Rueger et al. (2021) have already beautifully discussed this subject [10,14], and therefore, only points that could directly be addressed using megacolonies are considered below. Differences in anemonefish ecology, such as anemone host species, level of host specialization, capacities to move away from their hosts or not, etc., lead to various ecological constraints, which in turn create interspecific variations in social systems. In this way, comparing social behavior between species can help us to understand these variations’ proximate and ultimate causes. Megacolonies represent a model to study how social organization varies within a species, that is, the plasticity of the social behavior. Megacolonies are also a great opportunity to test the size-complexity hypothesis and assess social group transformation. What determines the ability of a species to form megacolonies? From our observations, both generalists (e.g., *A. clarkii, A. chrysopterus*) and specialists (e.g., *A. akallopisos, A. frenatus*) species can form bigger groups. As only little data is available, it is still unclear which species form or do not form megacolonies and what are the drivers of this alternative social structure. Elucidating answers to these questions would greatly help our understanding of what can drive such behavioral plasticity. We thus stress the need for more field observations.

Detailed studies of megacolonies could help address another exciting question: mating system plasticity. Could the strict monogamy usually observed in anemonefish be plastic when social and ecological constraints vary? Fishes display a great diversity of behavioral mating systems shaped by various environmental and behavioral parameters (densities and distribution, resources availability, level of parental care, territoriality) [44–46]. Plasticity of mating systems in fish is also quite common [47–50]. Thus, it could be expected that when habitat availability and group size increase, a switch toward polygamy could happen in anemonefishes, as in *D. aruanus* [51]. Occasional polygamy was observed in *A. clarkii*, with a male alternatively fertilizing clutches from two different females [33,36]. However, a detailed study by Fricke (1979) showed that monogamy was maintained, probably due to dominant males’ aggressive behavior toward smaller fish, which suppressed the maturation of testicular tissues [32]. Investigation of mating behavior in megacolonies could help gain insights into how conserved or plastic mating systems are in anemonefish. As females are known to be bigger and lay more eggs when living in larger hosts [52], estimating the lifetime reproductive success and parentage relations among colony members in megacolonies compared to “normal” colonies would also help in understanding how populations adapt to variable environmental conditions.

### Coexistence mechanisms

The use of megacolonies, particularly interspecific ones, as a model of coexistence, could help understand how species diversity is maintained, a crucial question in fundamental and applied sciences. Theoretical and empirical studies identify various ecological differences as the basis of species coexistence, and we now understand how species’ interactions with their environment can maintain species diversity [53–56]. For anemonefish, several studies have identified multiple mechanisms that sustain the coexistence of a large number of species [23,57,58]. Niche differentiation like resource partitioning by living in association with different anemone species or at different depths [23,57] is the main mechanism. But also cohabitation of different species occupying the same niche and habitat, and lottery, such as the chance to colonize vacant space, are at play [23,58]. However, the mechanisms promoting cohabitation are poorly known. Hattori (2002) suggested that differences in body sizes (big *A. clarkii* and small *A. perideraion*) are key to the cohabitation between those two species [59]. Coexistence could also vary depending on the life stage [60] as coexistence is sometimes observed only with juveniles [23] and host preference and mobility are known to depend on the development stage [61]. Reproductive interactions are also known to play a role in maintaining species diversity [62], which could be the case for anemonefish, given their particular mating system. We believe that interspecific megacolonies are very interesting models for investigating the diversity of mechanisms fostering species coexistence.

### Ultimate and proximate causes of aggressive behavior

In “normal” colonies, the dominant female and sub-dominant male are very aggressive and defend the colony against intruders, including divers or sharks [63]. In the megacolonies we observed, this behavior sometimes seemed to be either exacerbated (interspecific megacolonies in Taiwan) or lowered (intraspecific megacolony in Bora-Bora). In the second case, anemonefishes showed reduced aggressiveness among themselves and toward intruders (divers, *D. trimaculatus*). This would need further investigation and proper quantification and could offer a very interesting entry point to understand better the molecular mechanisms controlling aggressive behavior in anemonefishes [64,65]. It is tempting to relate the lower level of aggressiveness to an often-observed behavior in lab-reared juveniles (1-2 cm). Indeed, juvenile anemonefish are less aggressive when maintained at high densities than when maintained at low densities allowing them to establish a territory [66]. Whether having a well-defined territory to defend is a signal that promotes aggressive behavior, is an interesting hypothesis to test.

### An anemone’s perspective

Besides representing an exciting model to study anemonefish sociality and coexistence, megacolonies could also provide an excellent opportunity to investigate host anemones’ reproductive strategies. Host anemones have complex reproductive biology that remains generally poorly understood. They are gonochoric animals capable of sexual and asexual reproduction [67], but the extent of each reproductive strategy and the conditions inducing one or the other are unknown. Likewise, pelagic larval duration and settlement mechanisms are understudied [68].

It could be hypothesized that large anemone carpets could be formed by clones of the same or a few well-adapted individuals when environmental conditions are highly favorable [69]. In contrast, sexual reproduction would be favored when environmental conditions become less suitable, and dispersion to novel environments becomes a better option. We observed large clusters of *E. quadricolor*, for which asexual reproduction via longitudinal fission is well known [70,71], but also of *H. magnifica*, for which evidence of clonal reproduction is rare [72,73]. The formation of large clusters could then result from a higher larval settlement. Field monitoring and genetic surveys [74] of anemones clusters could help answer the following questions: are clusters composed of different genotypes, and if so, which factors are triggering higher larval recruitment (hydrodynamic, substrate type, light, conspecific density, etc.)? Or are they composed of clones, and are these anemone species more prone to clonal duplication, or do some environmental conditions enhance clonality over sexual reproduction? Understanding these mysterious animals’ reproductive ecology is an exciting field of research and could also help implement better conservation and management measures. Indeed, giant sea anemones are particularly targeted by fisheries for the aquarium trade and are sensitive to environmental disturbances. Their populations can withstand only slight pressure and need extended recovery times, and as anemonefish are obligate symbionts, the same applies to them [75].

### An overlooked concept

Most of the recent work on anemonefish’s social behavior has been done on *A. percula* (reviewed in [14,15]) and, to a lesser extent, *A. ocellaris* [76] but little is known about other anemonefish species. The only work performed on bigger groups and different species is now over 30 years old [33–36] and seems to have been overlooked or perhaps even forgotten by the scientific community. However, a detailed investigation of megacolonies would greatly benefit our scientific understanding of social group evolution, coexistence mechanism, aggressive behavior mechanisms, and even anemones’ ecology. We strongly encourage future research to consider this alternative social organization as a model worthy of more investigation. Results from such studies could also benefit the field of conservation biology. Indeed, variations in environmental conditions are known to affect social interactions. Therefore, as the frequency and intensity of environmental disturbances keep increasing [77], it seems urgent to understand the plasticity of intra- and inter-specific interactions in the face of changing environments to implement adequate conservation and management measures.

## Acknowledgments

The authors would like to thank Pr. Timothy Ravasi as well as Chanyoung Kim for their help in the field.

## Supporting information

**S1 Fig**. Bora-Bora megacolony

**S2 Fig**. Kagoshima megalony (video)

**S3 Fig**. Chinen and Oura bay megacolonies

**S4 Fig**. Kueishan megacolony (video)

**S5 Fig**. Atsuta megacolony (video)

**S6 Fig**. Aggressive behavior between juveniles (video)

